# Macrophage subpopulation identity in *Drosophila* is modulated by apoptotic cell clearance and related signalling pathways

**DOI:** 10.1101/2023.09.14.557767

**Authors:** Elliot C. Brooks, Martin P. Zeidler, Albert C. M. Ong, Iwan R. Evans

**Affiliations:** School of Medicine and Population Health and the Bateson Centre, University of Sheffield, UK; Department of Bioscience and the Bateson Centre, University of Sheffield, UK

## Abstract

In *Drosophila* blood, plasmatocytes of the haemocyte lineage represent the functional equivalent of vertebrate macrophages and have become an established *in vivo* model with which to study macrophage function and behaviour. However, the use of plasmatocytes as a macrophage model has been limited by a historical perspective that plasmatocytes represent a homogenous population of cells, in contrast to the high levels of heterogeneity of vertebrate macrophages. Recently, a number of groups have reported transcriptomic approaches which suggest the existence of plasmatocyte heterogeneity, while we identified enhancer elements that identify subpopulations of plasmatocytes which exhibit potentially pro-inflammatory behaviours, suggesting conservation of plasmatocyte heterogeneity in *Drosophila*. These plasmatocyte subpopulations exhibit enhanced responses to wounds and decreased rates of efferocytosis when compared to the overall plasmatocyte population. Interestingly, increasing the phagocytic requirement placed upon plasmatocytes is sufficient to decrease the size of these plasmatocyte subpopulations in the embryo. However, the mechanistic basis for this response was unclear. Here, we examine how plasmatocyte subpopulations are modulated by apoptotic cell clearance (efferocytosis) demands and associated signalling pathways. We show that loss of the phosphatidylserine receptor Simu prevents an increased phagocytic burden from modulating specific subpopulation cells, while blocking other apoptotic cell receptors revealed no such rescue. This suggests that Simu-dependent efferocytosis is specifically involved in determining fate of particular subpopulations. Supportive of our original finding, mutations in *amo* (the *Drosophila* homolog of *PKD2*), a calcium-permeable channel which operates downstream of Simu, phenocopy *simu* mutants. Furthermore, we show that Amo is involved in the acidification of the apoptotic cell-containing phagosomes, suggesting that this reduction in pH may be associated with macrophage reprogramming. Additionally, our results also identify Ecdysone receptor signalling, a pathway related to control of cell death during developmental transitions, as a controller of plasmatocyte subpopulation identity. Overall, these results identify fundamental pathways involved in the specification of plasmatocyte subpopulations and so further validate *Drosophila* plasmatocytes as a heterogeneous population of macrophage-like cells within this important developmental and immune model.

## Introduction

Macrophages are highly phagocytic cells of the vertebrate innate immune system, which are responsible for tissue homeostasis, fighting infection and removing apoptotic cells (Gordon & Plüddemann, 2017). Heterogeneity of the vertebrate macrophage is a fundamental component of the immune system, allowing these cells to respond to a variety of stimuli in a wide range of environments through differentiation into a range of tissue resident cell types and an ability to adopt various activation states, termed macrophage polarisation (Murray, 2017; Orecchioni et al., 2019). These activation states range from pro-inflammatory states associated with microbicidal activities (M1-like), to anti-inflammatory states, associated with wound healing and apoptotic cell clearance (M2-like), with this spectrum of activation states regulated in response to immediate environmental challenges (Hume, 2015). Aberrant macrophage polarisation has been implicated in numerous chronic inflammatory conditions, such as Chronic Obstructive Pulmonary Disease (COPD) and atherosclerosis, which are associated with increased M2-like polarisation and M1-like polarisation, respectively (Cornwell et al., 2018; de Gaetano et al., 2016; Shaykhiev et al., 2009). Though numerous experimental models have been exploited to facilitate our understanding of these fundamental processes, these often rely on *ex vivo* approaches that do not fully reproduce the temporal and spatial dynamics of *in vivo* biological systems. As such, low complexity *in vivo* models to study macrophage heterogeneity *in situ* have the potential to provide unique biological insights.

The fruit fly *Drosophila melanogaster* possesses an innate immune system comprising three lineages of haemocytes, specified via Serpent (Srp), the fly orthologue of the GATA transcription factors involved in vertebrate hematopoiesis (Lebestky et al., 2000). The plasmatocyte lineage is the dominant blood cell throughout normal development and represents the functional equivalent of vertebrate macrophages. Plasmatocytes mediate the same essential functions as macrophages, responding to wounds, fighting infection and removing apoptotic cells (efferocytosis) (Wood & Martin, 2017). Efferocytosis, has been particularly well studied in the fly, and multiple apoptotic cell receptors have been characterised including Simu and Draper (both CED1 family members), and Croquemort, which is homologous to the CD36 scavenger receptor expressed on human macrophages (Franc et al., 1996; Freeman et al., 2003; Kurant et al., 2008). Mutations in these receptors prevent efficient identification of dying cells, resulting in a build-up of uncleared apoptotic corpses *in vivo*, the persistence of which disrupts other plasmatocyte functions, such as migration and wound responses (Evans et al., 2015; Roddie et al., 2019). However, despite the undoubted similarities between plasmatocytes and macrophages, until recently there was limited evidence to suggest that plasmatocytes were as functionally or molecularly diverse as their vertebrate counterparts.

Recent studies into plasmatocyte behaviour *in vivo* reveal that these cells do not behave in a uniform manner. Following their dispersal across the embryo, plasmatocytes surrounding the ventral nerve cord appear to move randomly as if they are no longer migrating towards chemotactic cues. Imaging this random movement in stage 15 embryos revealed a wide range in migration speeds, suggesting some plasmatocytes may be developmentally programmed to have enhanced motility capabilities. Similarly, imaging plasmatocyte movements in the vicinity of a sterile wound indicated that some plasmatocytes that that exhibit a rapid and robust migratory response, while others at similar distances from the wound site fail to respond (Coates et al., 2021; Roddie et al., 2019). There is also remarkable variability in the number of apoptotic corpses phagocytosed by plasmatocytes (Coates et al., 2021; Raymond et al., 2022; Roddie et al., 2019). These data hint that the overall plasmatocyte population is made up of subpopulations, each of which exhibit distinct innate immune behaviours. Consistent with this cellular diversity, transcriptional profiling studies using single cell RNA sequencing (scRNAseq) approaches confirm the existence of molecularly-defined plasmatocyte clusters in larvae (Cattenoz et al., 2020; Cho et al., 2020; Fu et al., 2020; Tattikota et al., 2020), while reporter studies indicate heterogeneity across the *Drosophila* lifecourse (Coates et al., 2021).

In addition to a diversity of activities at any one developmental stage, the behaviour of *Drosophila* plasmatocytes also changes markedly throughout the life cycle of the fly. During embryogenesis, plasmatocytes are highly migratory as they disperse around the embryo, shaping tissues and organs via deposition of extracellular matrix (Matsubayashi et al., 2017) and phagocytosis of apoptotic cells (Page & Olofsson, 2008). By contrast, plasmatocytes are largely sessile during larval stages, adhering to the body wall where they proliferate under the control of Activin-α released from nearby neurons (Leitão & Sucena, 2015; Makhijani et al., 2017). At the onset of metamorphosis, plasmatocyte behaviour changes once more, as these cells are reprogrammed to become more highly phagocytic and migratory to deal with the high levels of cell death associated with the tissue remodelling during this developmental stage (Regan et al., 2013). These alterations in plasmatocyte behaviour are influenced by the action of the steroid hormone 20-hydroxyecdysone (hereafter referred to as ecdysone), itself central to progression through these developmental stages which require significant tissue remodelling and apoptosis. Whether these changes in behaviour are linked to changes in subpopulation identity is unknown. Further examples of the importance of ecdysone in behavioural transition points is illustrated by the fact that embryonic plasmatocytes expressing a dominant-negative isoform of the ecdysone receptor (EcR) fail to mount effective responses to infection (Tan et al., 2014), while increasing ecdysone levels acting via the ecdysone receptor B1 stimulate plasmatocytes to become highly motile and phagocytic during pupation (Jiang et al., 1997; Sampson et al., 2013; Zirin et al., 2013). Thus, plasmatocytes exhibit plasticity, with their behaviours changing across the lifecourse, according to the developmental stage, with the action of ecdysone implicated in contributing to these changes.

Following up *in vivo* evidence hinting at plasmatocyte heterogeneity and plasticity, we recently exploited the Vienna Tiling (VT) array library of non-coding enhancer elements (Kvon et al., 2014) and identified enhancers active in a subset of plasmatocytes (Coates et al., 2021). This approach revealed the existence of functionally-distinct subpopulations that were associated with enhanced wound responses and decreased rates of apoptotic cell clearance compared to the overall plasmatocyte population, behaviours which are typical of pro-inflammatory macrophages. These subpopulations are developmentally regulated, with relatively high numbers in embryonic stages preceding a drastic decrease in larval stages, before subpopulations ultimately re-emerge at the onset of metamorphosis and persist into adulthood (Coates et al., 2021), patterns consistent with the changes in plasmatocyte behaviour seen during development, including those regulated by ecdysone signalling.

The plasticity of *Drosophila* plasmatocytes can also be observed by increasing the apoptotic challenge they face *in vivo*. During embryogenesis, the other major cell-type involved in efferocytosis are glial cells, specified via the transcription factor *repo* (Halter et al., 1995; Kurant et al., 2008; Sonnenfeld & Jacobs, 1995). Loss of glial specification in *repo* null embryos results in increased levels of uncleared apoptotic cells – a change previously shown to impair plasmatocyte migration and wound responses (Armitage et al., 2020). Interestingly, the plasmatocyte subpopulations studied to date exhibit a significant decrease in their relative numbers in a *repo* mutant background (Coates et al., 2021); this suggests that the high- apoptotic environment seen in *repo* mutants may either prevent plasmatocytes acquiring pro-inflammatory subpopulation identities or alternatively drive exit from those subpopulations.

The fact that putative pro-inflammatory macrophage subpopulations decrease in the presence of high levels of uncleared apoptotic cells (Coates et al., 2021) suggests that signalling downstream of apoptotic cell receptors may influence subpopulation fate. Here, we show that Simu, a receptor for apoptotic cells, mediates decreases in the numbers of specific plasmatocyte subpopulation cells on exposure to enhanced levels of apoptosis. Furthermore, mutations affecting the calcium-permeable cation channel Amo, which regulates calcium homeostasis downstream of phagocytosis (Van Goethem et al., 2012), phenocopy results seen in *simu* mutants. This is consistent with a model whereby Amo functions downstream of Simu to facilitate plasmatocyte reprograming to alternative fates. We also identify a defect in phagosome acidification in *amo* mutants, suggesting that this process may be important for the removal of plasmatocytes from specific subpopulations. Finally, we demonstrate a requirement for ecdysone signalling in the establishment of subpopulation identity, both in the embryo and pupa. Overall, these findings further reinforce the utility of *Drosophila* plasmatocytes as a robust macrophage model – with cells clearly exhibiting heterogeneity, plasticity, and the apparent ability to switch activation states in response to different environmental challenges.

## Methods

### Fly genetics and reagents

Stocks of *Drosophila* were kept at 25°C on standard cornmeal/molasses agar. To collect embryos, at least 20 male and 20 female flies were placed in a 100mL beaker which was capped with a 5cm apple juice agar plate supplemented with a small amount of yeast paste (50% in dH2O), secured with an elastic band and incubated at 22°C for embryo collections. For pupal experiments, crosses were kept at 25°C and white pre-pupae were collected during 30-minute windows and aged at 25°C. All transgenes and mutations were crossed into a *w^1118^* background.

The following *Drosophila* drivers and constructs were used: *crq-GAL4* (Stramer et al., 2005), *srpHemo-GAL4* (Brückner et al., 2004), *hml(Δ)-GAL4* (Sinenko & Mathey-Prevot, 2004), *VT17559-GAL4, VT32897-GAL4, VT57089-GAL4, VT62766- GAL4* (Coates et al., 2021; Kvon et al., 2014). When exploiting the split GAL4 system (Pfeiffer et al., 2010), the GAL4 activation domain (AD) was expressed via *srpHemo-AD*, while the GAL4 DNA binding domain (DBD) was expressed via *srpHemo-DBD*, *VT17559-DBD*, *VT32897-DBD, VT57089-DBD* and *VT62766-DBD* (Coates et al., 2021). The following UAS lines were used: *UAS-stinger* (Barolo et al., 2000), *UAS-eGFP* (Bloomington *Drosophila* Stock Centre), *UAS-Draper-II* (Logan et al., 2012), and *UAS-EcR.B1^ΔC655^* (Cherbas et al., 2003). The following GAL4-independent reporter lines were used: *srpHemo-3x-mCherry*, *srpHemo-H2A-3x-mCherry* (Gyoergy et al., 2018), *srpHemo-GMA* (James Bloor, University of Kent), *VT17559-RFP, VT32897-RFP, VT57089-RFP* and *VT62766-RFP* (Coates et al., 2021). The following *Drosophila* mutants were used: *amo^1^* (Watnick et al., 2003), *simu^2^* (Kurant et al., 2008), *repo^03702^* (Xiong et al., 1994) and *crq^ko^* (Guillou et al., 2016).

### Imaging of *Drosophila* embryos

All embryos were dechorionated in bleach (Evans et al., 2010) prior to being mounted ventral-side-up on double-sided tape (Scotch) in a minimal volume of Voltalef oil (VWR). All imaging of embryos was carried out on an UltraView Spinning Disk System (Perkin Elmer) using a 40x UplanSApo oil immersion objective lens (NA 1.3). Embryos were imaged to a depth of 20µm, with z-slices spaced every 1µm. For Lysotracker experiments requiring staining of live embryos, stage 15 dechorionated embryos were selected and transferred to a mixture of peroxide-free heptane (VWR) and 10μM lysotracker red (Thermofisher) in PBS (Oxoid) in a glass vial, which was shaken in the dark for 30 minutes. Embryos were then transferred into a Watchmaker’s glass containing Halocarbon oil 700 (Sigma), before being mounted as described above.

Embryos requiring fixation and immunostaining were fixed and stained as previously described (Roddie et al., 2019). For Fascin staining, embryos were treated with a mouse anti-Fascin primary antibody (sn7c; Developmental Studies Hybridoma Bank; used at 1:500) and Alexa fluor 568 goat anti-mouse (A11031; Life Technologies; used at 1:200).

### Imaging of pupae

Pupa of the appropriate genotype were selected and aged to 48h after puparium formation (APF) at 25°C and attached to slides via double-sided tape. Pupae were carefully removed from their pupal cases and covered in a small volume of Voltalef oil. Stacks of 5 coverslips (22 x 22mm, thickness 1) were glued together with nail varnish and then placed either side of pupae. A coverslip (22 x 32mm, thickness 1) was then placed over the top of the pupae, in contact with the oil, supported by the coverslip stacks to prevent damage to pupae. Z-stacks were then taken of the thoracic regions of pupae using a Nikon spinning disk system (Nikon Eclipse Ti2 microscope with a CSU-W1 Okagawa confocal scanner unit and Photometrics Prime 95B 22mm camera; 20X Plan Apo/0.75 objective lens, GFP and RFP filters, 2μm between slices).

### Image analysis and statistical analyses

All images were converted to .tiff format, despeckled to reduce background noise, and blinded prior to analysis, which was performed using Fiji/ImageJ (Schindelin et al., 2012). To work out the relative number of plasmatocytes within subpopulations in embryos, z-stacks were converted into a maximum intensity projection in Fiji. The total number of plasmatocytes, labelled with pan-plasmatocyte reporters (such as *srpHemo-3x-mCherry*, *crq-GAL4,UAS-GFP;srpHemo-GAL4,UAS-GFP* or anti-Fascin staining), was counted. Numbers of subpopulation plasmatocytes were quantified from images of cells labelled using *VT-GAL4* lines (Kvon et al., 2014), or *srpHemo-AD* in concert with *VT-DBD* transgenes (Coates et al., 2021) to drive UAS reporters, or via GAL4-independent *VT-RFP* lines (Coates et al., 2021). The proportion of plasmatocytes within a given subpopulation was then expressed as a percentage of the overall population.

To quantify phagosome acidification, the number of phagosomes of the 5 most-ventral plasmatocytes were selected using the multi-point selection tool and the number of these phagosomes which overlapped with lysotracker red staining were then counted to work out a percentage of phagosome acidification.

For quantification of subpopulations within the pupal thorax (at 48h APF), maximum projections corresponding to 10 z-slices were assembled starting from the z-position in which the most superficial plasmatocytes were visible (labelled via *hml(Δ)-GAL4,UAS-GFP*), moving deeper into the pupa. All images in the dataset underwent identical contrast adjustment and were blinded ahead of analysis. Cells were then scored as *hml*-positive only or double-positive for *hml* and *VT62766* on the basis of the presence/absence of visible fluorescence in the GFP (*hml(Δ)-GAL4,UAS-GFP*) and RFP (*VT62766-RFP*) filter channels. Cells obscured by the bounding vitelline membrane (“rings” in the RFP channel in Fig. 7) were not assessed. The multi-point tool in Fiji was used to keep track of cells that had been assessed. To normalise total numbers of *hml*-positive cells within the thoracic region, the area bounded by the vitelline membrane was measured using the polygon selection tool in Fiji. As an additional means of quantification, the same maximum projections were cropped to the region of interest (demarcated by the vitelline membrane). The green channel was then manually thresholded to create a mask corresponding to plasmatocyte localisation. This was then used to measure total fluorescence (integrated density) within *hml*-positive plasmatocytes for GFP and RFP channels to quantify reporter activity.

Statistical analysis was conducted in GraphPad Prism. Mann-Whitney and Student’s unpaired t-tests were used to compare non-parametric and parametric data, respectively. Where greater than two means were to be compared a one-way ANOVA with Dunnett’s post-test was used (parametric data).

## Results

### Loss of Simu, an apoptotic cell receptor, drives expansion of specific plasmatocyte subpopulations

We previously described how numbers of haemocytes in subpopulations defined by the *VT17559*, *VT32897*, *VT57089* and *VT62766* enhancers are reduced in a genetic background containing excess apoptotic cells (*repo* mutants; Armitage et al., 2020; Coates et al., 2021). To explore the mechanistic basis for this effect, we examined the effect of loss of *simu* (*six microns under*); *simu* encodes a receptor used by macrophages to recognise phosphatidylserine present on the surface of apoptotic cells (Kurant et al., 2008). Given that the absence of *simu* in embryos has previously been shown to increase levels of uncleared apoptotic cells (Kurant et al., 2008; Roddie et al., 2019), we determined the proportion of macrophage subpopulations in stage 15 embryos lacking *simu* versus control embryos (Fig. 1). Subpopulation plasmatocytes were labelled via the split GAL4 system (Pfeiffer et al., 2010) driving *UAS-GFP* specifically in cells with overlapping expression of *serpent* (*srpHemo-AD*) and the subtype-specific VT enhancer (*VT-DBD*) activity (Coates et al., 2021), while the total plasmatocyte population was independently labelled via a GAL4-independent reporter (*srpHemo-3x-mCherry*; Gyoergy et al., 2018; labelled in magenta in Fig. 1).

**Figure 1.**
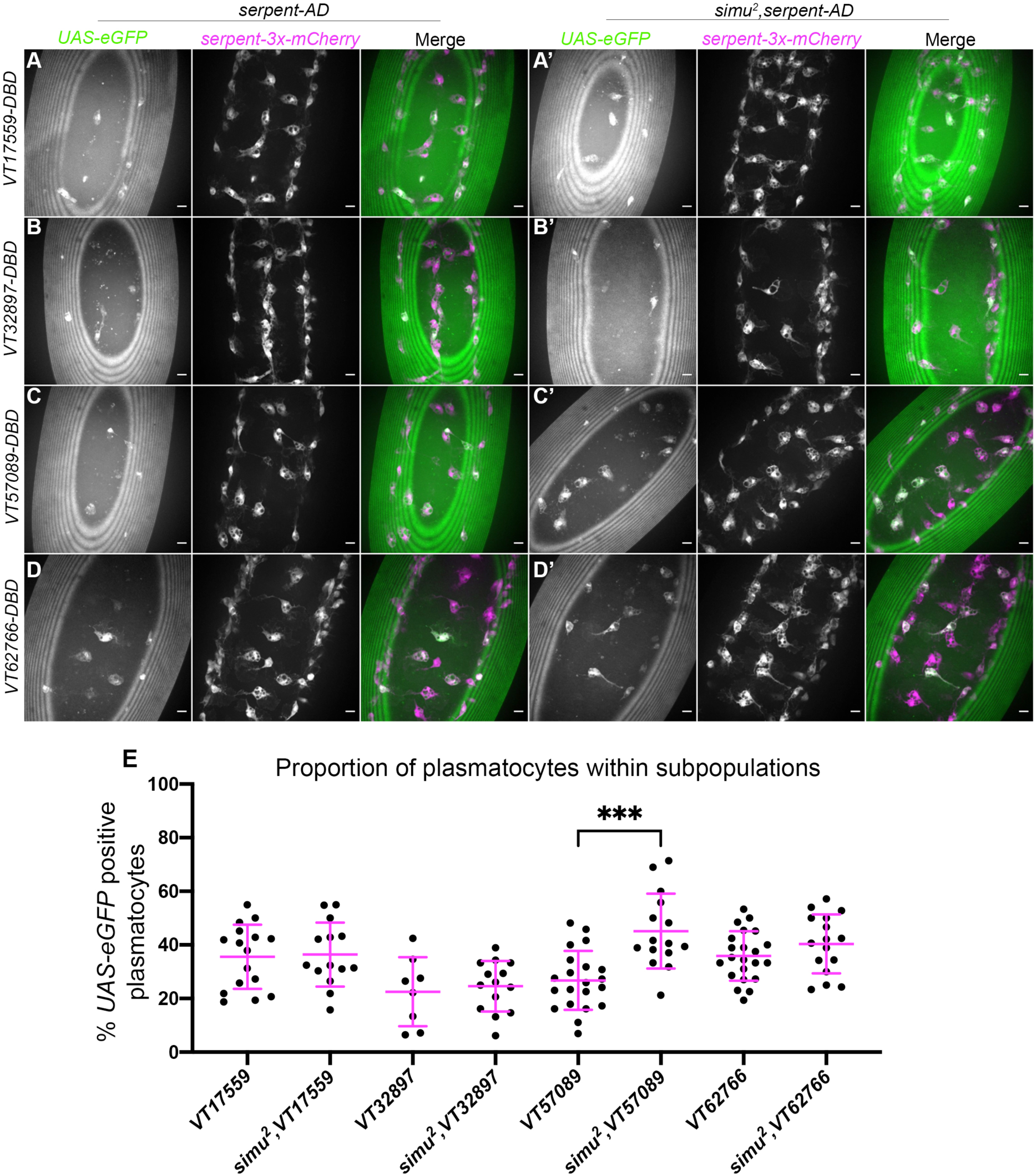
Overexposure of plasmatocytes to apoptotic cells via loss of Simu is not sufficient to cause a decrease in plasmatocyte subpopulations. (A-D’) representative images of the ventral midline of control (A-D) and *simu^2^* (A’-D’) embryos at stage 15. *UAS-eGFP* shows plasmatocytes labelled via split-*GAL4* (green in merge), while *srpHemo-3x-mCherry* labels every plasmatocytes (magenta in merge). Anterior is up in all images, scale bars denote 10µm. (E) scatterplot showing proportion of plasmatocytes within subpopulations in control and *simu^2^* embryos. *n*=16, 14, 8, 14, 21, 15, 22 and 16, respectively. Only significantly different results shown on the graph (p=0.0003), all statistical comparisons carried out via non-paired t-tests. *** P<0.001.

In contrast to *repo* mutants, where an increased apoptotic challenge correlates with a decrease in numbers of subpopulation plasmatocytes (Coates et al., 2021), no change was observed for the *VT17559*, *VT32897* and *VT62766* subpopulations in a *simu* mutant background (Fig. 1A-E), while numbers of *VT57089*-labelled cells actually increased in *simu* mutants compared to controls (Fig. 1C, E). This suggests that Simu normally antagonises acquisition or maintenance of the *VT57089* fate. These results suggest that the relative decreases in subpopulation cells seen in *repo* mutants (Coates et al., 2021) may depend upon the presence of Simu on the surface of plasmatocytes – a receptor required for effective recognition and phagocytosis of apoptotic cells (Kurant et al., 2008).

### Simu mediates antagonism of specific subpopulation fates in the presence of large amounts of apoptosis

The contrasting results seen between *repo* and *simu* mutant embryos indicate that increased levels of uncleared apoptotic cells may not be sufficient to mediate decreases in subpopulation numbers – instead, specific interactions between apoptotic cells and plasmatocytes may be required for phenotypic switches. To test whether effective recognition and/or engulfment of apoptotic cells is responsible for mediating the reduction in subpopulation plasmatocytes observed in *repo* mutants, *simu;repo* double mutants were generated and compared to controls (as well as *simu* and *repo* single mutants). Due to the genetics involved in this experiment, subpopulation plasmatocytes were labelled via *VT-GAL4* (as opposed to split *VT- GAL4*) driving expression of *UAS-stinger* (Fig. 2A-D).

**Figure 2.**
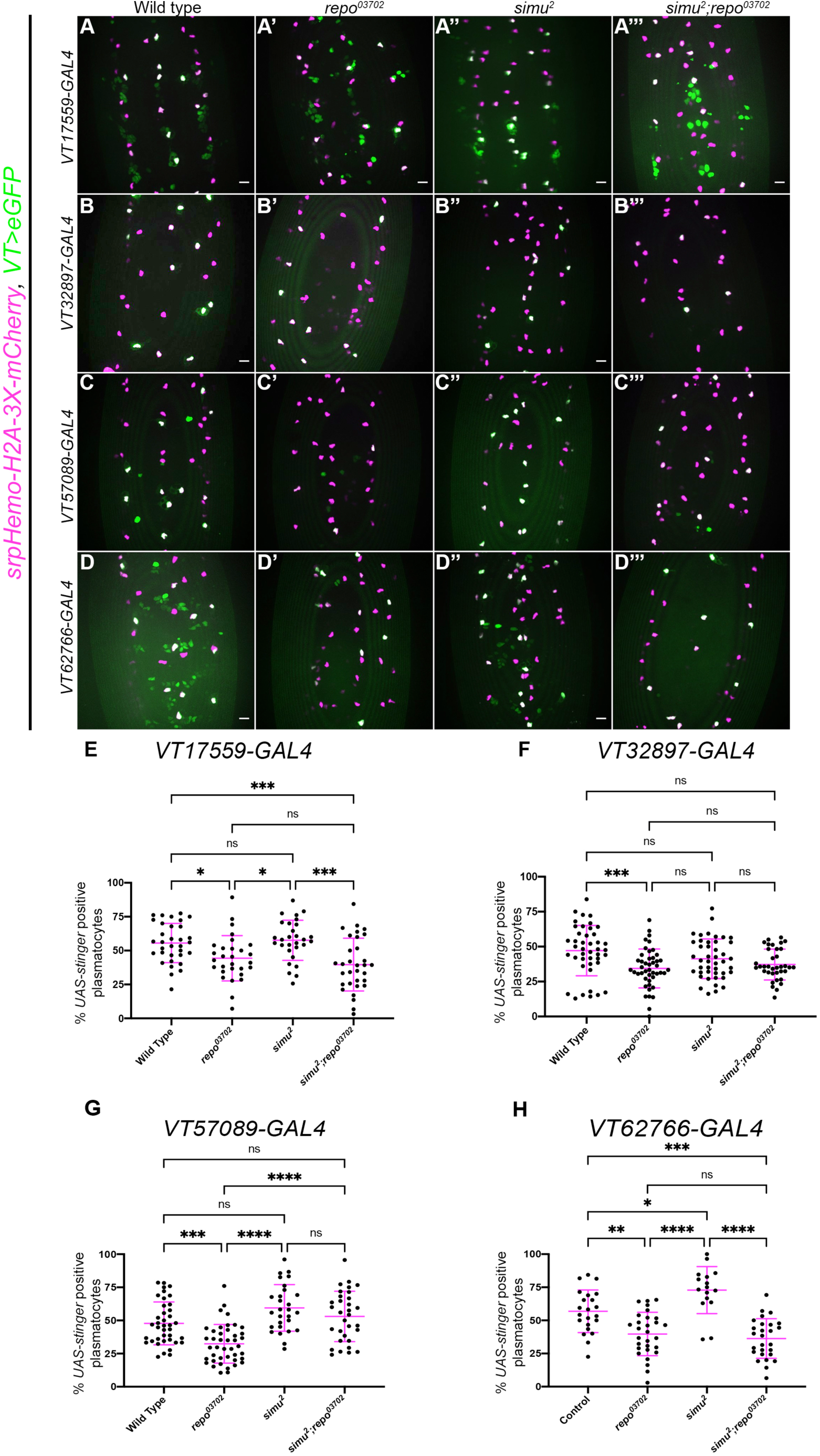
Simu-dependent efferocytosis is required for the decrease in the *VT57089* subpopulation seen in *repo* mutants. (A-D’’’) representative maximum projection images of the ventral midline of control (A-D), *repo^03702^* single mutants (A’-D’), *simu^2^* single mutant (A”-D”) and *simu^2^;repo^03702^* double mutant (D’”-D’”) embryos at stage 15. *UAS-stinger* shows subpopulation plasmatocytes labelled via *VT*- *GAL4* (green in merge), while *srpHemo-H2A-3x-mCherry* labels every plasmatocytes (magenta in merge). Anterior is up in all images, scale bars denote 10µm. (E) scatterplot showing proportion of plasmatocytes within the *VT17559* subpopulation. *n*=33, 29, 29 and 32 respectively. (F) scatterplot showing proportion of plasmatocytes within the *VT32897* subpopulation. *n*=44, 45, 43 and 36 respectively. (G) scatterplot showing proportion of plasmatocytes within the *VT57089* subpopulation. *n*=39, 42, 28 and 32 respectively. (H) scatterplot showing proportion of plasmatocytes within the *VT62766* subpopulation. *n*=22, 29, 16 and 27 respectively Statistical analyses carried out via one-way ANOVA with Dunnett’s multiple comparison test. *, **, *** and **** represent p<0.05, p<0.01, p<0.001 and p<0.0001, respectively.

As we have previously shown (Coates et al., 2021), fewer plasmatocytes were present in all subpopulations examined in the presence of *repo* mutations alone (Fig. 2E-H). The loss of *simu* in addition to *repo* (*simu;repo* double mutants) was unable to rescue relative plasmatocyte subpopulation numbers to control levels for the *VT17559* and *VT62766* reporters (Fig. 2E, H); there is considerable variability in subpopulation numbers and results for the *VT32897* reporter were not statistically significant (Fig. 2F), though the trends were consistent with those observed for *VT17559* and *VT62766* (Fig. 2E, H). This variability potentially stems, in part, from the stochastic nature of contact between apoptotic corpses and plasmatocytes in the embryo. In contrast, numbers of cells labelled via the *VT57089* reporter were completely rescued to control levels in *simu;repo* double mutants (Fig. 2G). This implies that Simu-dependent efferocytosis, or an effector downstream of Simu-dependent recognition of apoptotic cells, may mediate the apparent shift out of the *VT57089* subpopulation seen in the presence of large numbers of apoptotic cells. Furthermore, this suggests distinct mechanisms may control plasmatocyte identity from subpopulation to subpopulation in response to the high apoptotic challenge presented to plasmatocytes (as assayed via *repo* loss-of-function).

### Other apoptotic cell clearance receptors do not appear to contribute to subpopulation identity

In addition to *simu*, *croquemort* (*crq*) and *draper* (*drpr*) encode two further apoptotic cell clearance receptors expressed on the surface of embryonic *Drosophila* plasmatocytes (Franc et al., 1996; Manaka et al., 2004). Given the role of *simu*, we next examined the role of Crq and Drpr in regulation of subpopulation identity following challenge with large numbers of apoptotic cells (a *repo* mutant background).

A loss-of-function *crq* allele (*crq^ko^*; Guillou et al., 2016) was used alongside *repo* mutations to investigate whether Crq is required for the decreased numbers of subpopulation plasmatocytes seen in *repo* mutants. Subpopulation plasmatocytes were labelled via *VT-GAL4* driving *UAS-stinger*, while all plasmatocytes were labelled via immunostaining for Fascin, an Actin-bundling protein highly enriched in *Drosophila* plasmatocytes (Zanet et al., 2009; Fig. 3A- D). This approach revealed no differences in the proportion of plasmatocytes found within each subpopulation when comparing *repo* single mutants to *crq;repo* double mutants (Fig. 3E). This suggests that the decrease in subpopulation numbers seen in *repo* mutants is independent of Crq.

**Figure 3.**
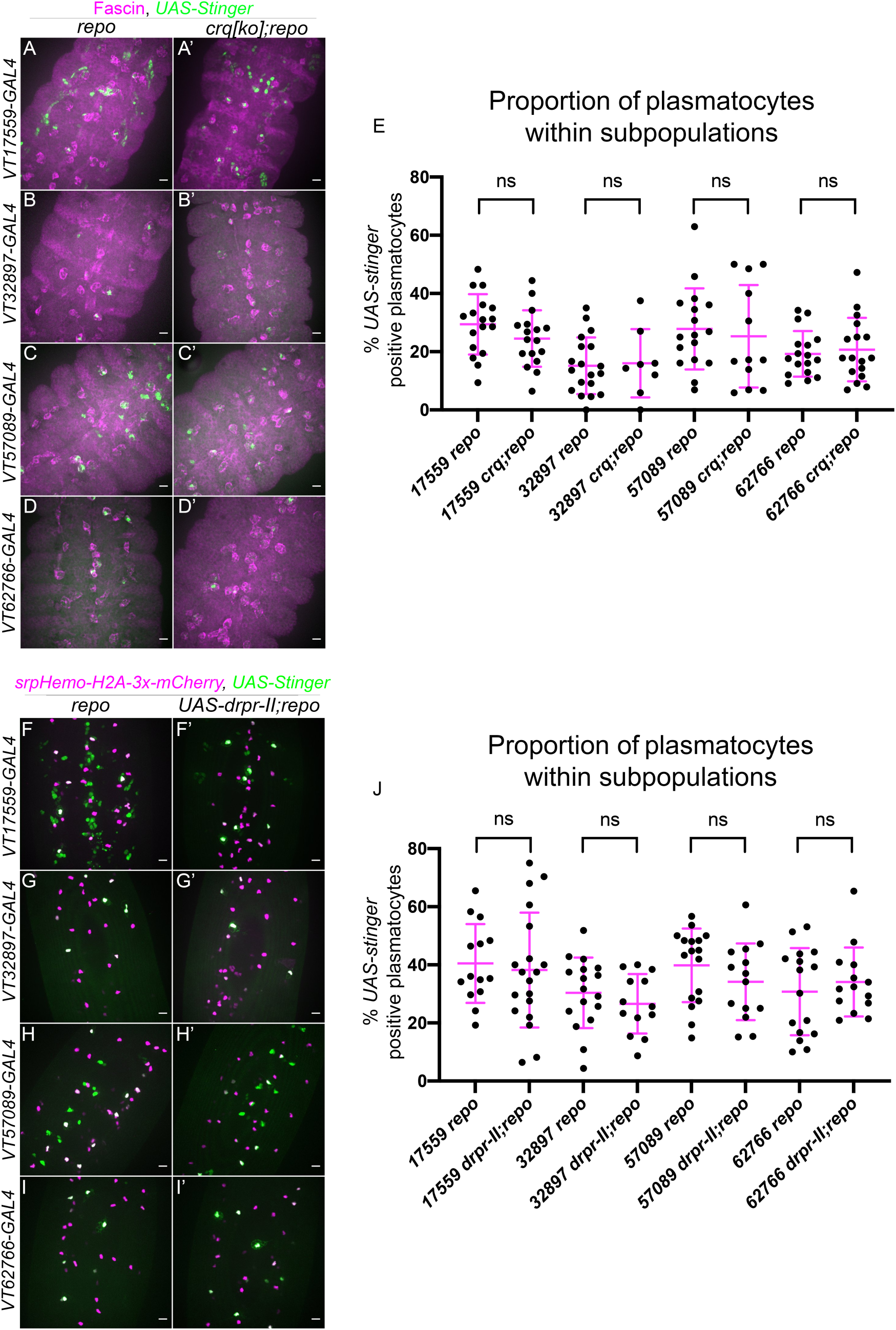
Croquemort and Draper do not affect plasmatocyte subpopulations. (A-D’) representative maximum projection images of the ventral midline of *repo* only single mutant embryos (A-D) and *crq;repo* double mutant embryos (A’-D’), at stage 15. Plasmatocytes labelled via Fascin staining (magenta) while subpopulation plasmatocytes are labelled via *VT- GAL4* driving expression of *UAS-Stinger* (green). Anterior is up in all images, scale bars denote 10µm. (E) scatterplot showing proportion of plasmatocytes within subpopulations. *n=* 18, 17, 20, 8, 17, 12, 17 and 17, respectively. Statistical analyses carried out via unpaired *t* tests. (F-I’) representative maximum projection images of the ventral midline of *repo* only single mutant embryos (F-I) and *repo* mutant embryos expressing *UAS-Drpr-II* specifically in plasmatocytes (F’-I’), at stage 15. All plasmatocytes labelled via *srpHemo-GAL4,UAS-GFP* and *crq-GAL4,UAS-GFP* while subpopulation plasmatocytes are labelled via *VT-RFP.* Anterior is up in all images, scale bars denote 10µm. (J) scatterplot showing proportion of plasmatocytes within subpopulations. *n*=14, 19, 17, 13, 17, 14, 16 and 14, respectively. Statistical analyses in (E and J) carried out via unpaired *t* tests.

To investigate the involvement of Drpr in modulating plasmatocyte subpopulations, an inhibitory isoform of Drpr (*UAS-Drpr-II*; Logan et al., 2012) was expressed specifically in subpopulation plasmatocytes. This was driven using *VT-GAL4*, which simultaneously enabled expression of *UAS-Stinger* to label subpopulation cells; the overall plasmatocyte population was labelled via *srpHemo-H2A-3x-mCherry* (Fig. 3F-I). Similar to the use of *crq* mutants, expression of Drpr-II specifically in subpopulation cells in a *repo* mutant background did not impact subpopulation numbers when compared to *repo* mutants lacking *UAS-Drpr-II* expression (Fig. 3J). Taken together this suggests that neither Drpr nor Crq modulate subpopulation identity at embryonic stages, in contrast to Simu.

### Amo functions downstream of Simu to control identity of specific subpopulations

Our results so far have shown that Simu appears to be involved in shifting plasmatocytes out of the *VT57089* subpopulation, both in control backgrounds (Fig. 1E) and in response to the high apoptotic challenge presented by *repo* mutations (Fig. 2G). The calcium-permeable cation channel Amo, homologous to human *PKD2*, which is causative of Autosomal Dominant Polycystic Kidney Disease (ADPKD), has previously been shown to maintain calcium homeostasis downstream of Simu during later stages of efferocytosis in *Drosophila* (Van Goethem et al., 2012). We therefore used a loss-of-function *amo* allele (*amo^1^*; Watnick et al., 2003) to investigate whether Amo-dependent calcium homeostasis is involved in modulating the identity of *VT57089* subpopulation plasmatocytes in stage 15 embryos. As per Fig. 1C, *VT57089* subpopulation plasmatocytes were labelled using the split GAL4 system to drive expression of *UAS-eGFP*, while the overall plasmatocyte population was labelled via *srpHemo- 3x-mCherry* (Fig. 4A-B).

**Figure 4.**
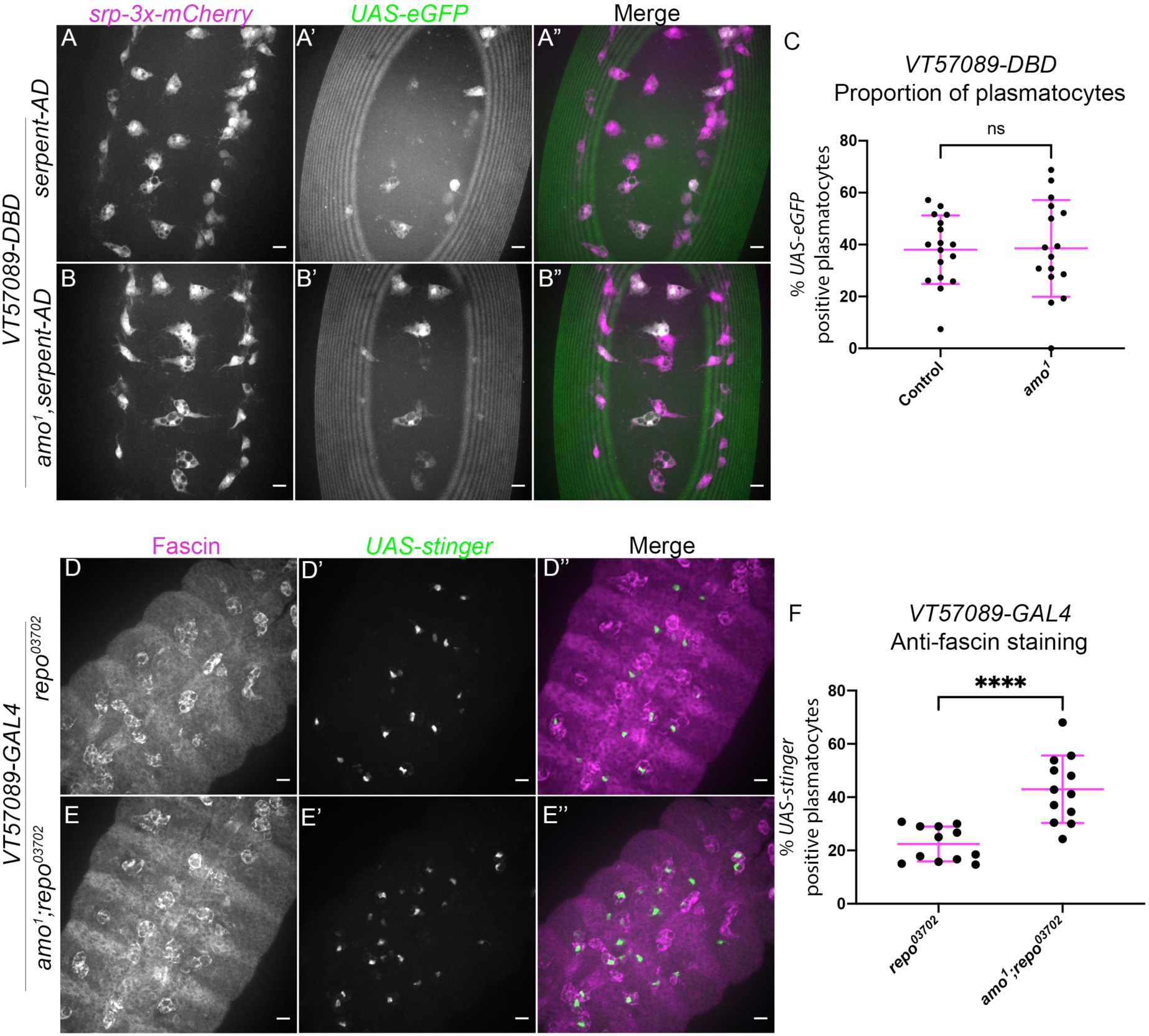
The calcium permeable cation channel Amo is required for the shift out of the *VT57089* subpopulation seen in *repo* mutants. (A-D”) representative images of the ventral midline of control (A-A”) and *amo^1^* (B-B”) embryos at stage 15. *srpHemo-3x-mCherry* labels every plasmatocyte (magenta in merge – A,B), while *VT57089* subpopulation plasmatocytes labelled via split *VT57089*-*GAL4* driving expression of *UAS-eGFP* (green in merge – A’,B’). Anterior is up in all images, scale bars denote 10µm. (C) scatterplot showing the proportion of plasmatocytes within the *VT57089* subpopulation in control and *amo^1^* embryos. *n*=17 and 16, respectively. (D-E”) representative maximum projection images of the ventral midline of *repo^03702^* single mutant (D-D”) and *amo^1^;repo^03702^* double mutant (E-E”) embryos at stage 15. Plasmatocytes have been labelled via Fascin staining (magenta in merge – D,E), with *VT57089* positive cells labelled via *UAS-stinger* (green in merge – D’,E’). Anterior is up in all images, scale bars denote 10µm. (F) scatterplot showing proportion of plasmatocytes within the *VT57089* subpopulation between *repo^03702^* single mutant and *amo^1^;repo^03702^* double mutant embryos based on anti-Fascin staining. *n*=12 and 13, respectively. Statistical analyses in (C and F) carried out via non-paired *t*-tests. Statistical analysis in (I) carried out via Mann-Whitney tests. **** represents p<0.0001.

Unlike *simu* mutants, no differences in the proportion of plasmatocytes within the *VT57089* subpopulation were observed when comparing wild-type embryos to *amo^1^* single mutants (Fig. 4C). These experiments were initially conducted in embryos with ‘normal’ – i.e., developmental levels – of apoptosis. We therefore next introduced *repo* mutations to address how loss of Amo impacted the *VT57089* subpopulation in response to an increased apoptotic challenge. Subpopulation plasmatocytes were labelled via *VT-GAL4* driving *UAS-stinger*, while anti-Fascin staining was again used to label the overall macrophage population to calculate the proportion of plasmatocytes within this subpopulation (Fig. 4D-E). Interestingly, these results phenocopied *simu* mutants, with a significantly higher proportion of *VT57089* plasmatocytes found in *amo^1^;repo* double mutants compared to *repo* only controls (Fig. 4F). This suggests that Amo-mediated calcium homeostasis is an important aspect of signalling downstream of Simu, which may prevent acquisition of *VT57089* identity, or mediate reprogramming of plasmatocytes away from this specific subpopulation.

To investigate the functional impact of *amo* mutations in the presence of elevated apoptotic challenge – i.e., in a *repo* mutant background (Armitage et al., 2020; Shklyar et al., 2014) – lysotracker staining was utilised to visualise defects in phagosome acidification. Phagosome acidification occurs during the later stages of efferocytosis, downstream of engulfment (Kinchen & Ravichandran, 2008), and is a calcium-dependent process (Westman et al., 2019). Thus, this process was an attractive pathway to investigate with respect to *amo* mutations, due to the calcium permeability of the Amo cation channel. All plasmatocytes were labelled via *crq-GAL4* driving expression of *UAS-eGFP* (Fig. 5A-C), and the number of acidified (lysotracker red positive) vacuoles per plasmatocyte was quantified. Lysotracker staining of *repo* single mutants and *amo;repo* double mutants revealed a significant reduction in the total number of acidified phagosomes present within in *amo*;*repo* double mutant plasmatocytes compared to *repo* only controls (Fig. 5D). Furthermore, the proportion of phagosomes that were acidified was also significantly lower in *amo;repo* double mutants (Fig. 5E). These results suggest that Amo is either required for phagosomal acidification or plays an upstream role in that process during efferocytosis. Defects in acidification may interfere with specification of plasmatocytes as *VT57089* subpopulation cells, alternatively, this process could lead to plasmatocytes exiting this subpopulation fate. Overall, we have shown that Simu negatively regulates *Drosophila* plasmatocyte subpopulation identity. *Amo* also appears required, with its role potentially linking effective phagosome acidification during the later stages of efferocytosis to regulation of cell identity.

**Figure 5.**
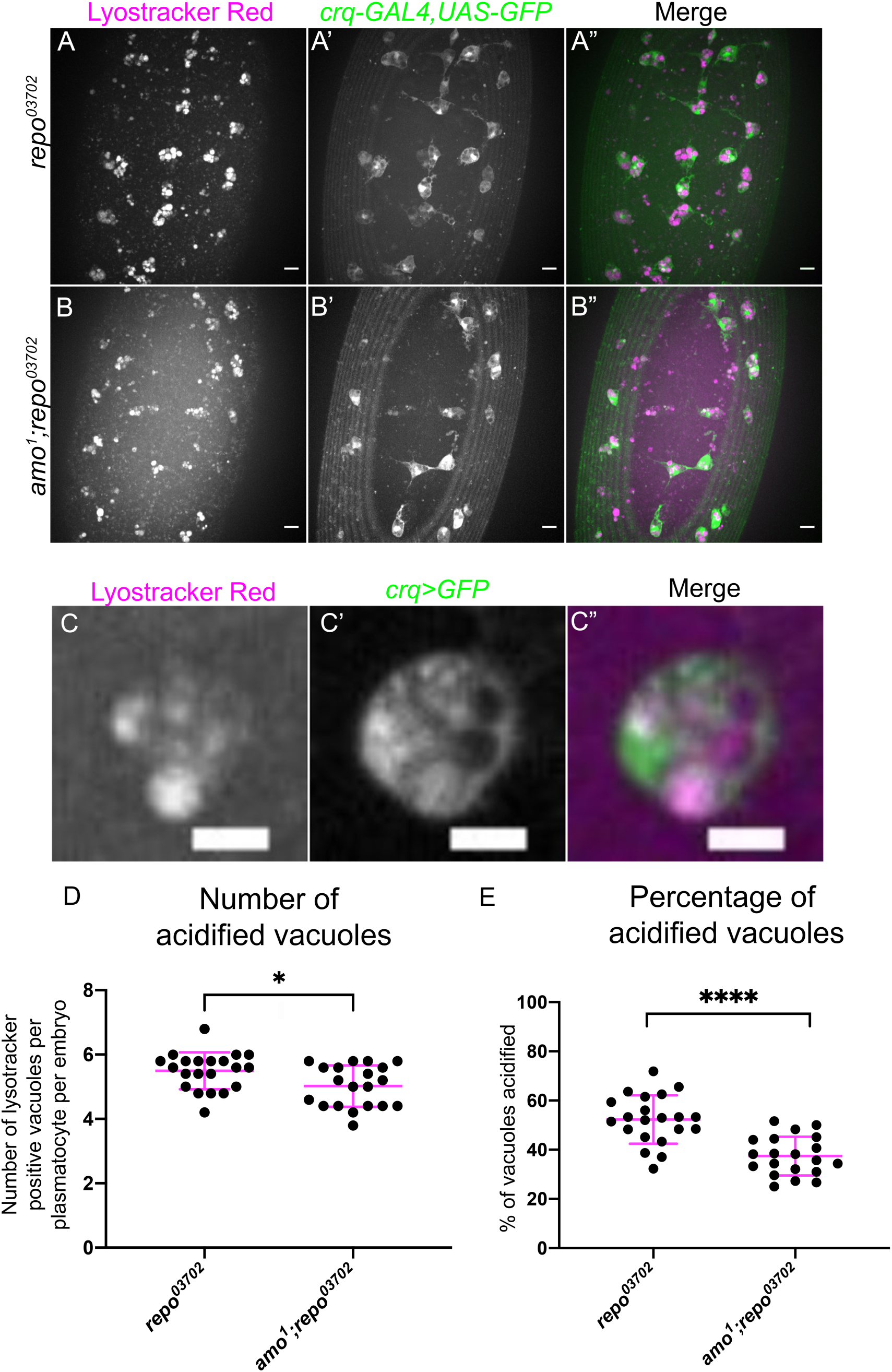
Amo is required for effective phagosome acidification. (A-B”) representative maximum projection images of the ventral midline of *repo^03702^* single mutant (A-A”) and *amo^1^;repo^03702^* double mutants (B-B”), embryos at stage 15. Lysotracker red shows acidified phagosomes (A-B), while all plasmatocytes are labelled via *crq-GAL4,UAS-eGFP* (A’-B’) . Anterior is up in all images, scale bars denote 10µm. (C-C”) zoom of a representative plasmatocyte showing both acidified and non-acidified phagosomes. Scale bar denotes 5µm. (D) scatterplot showing average number of lysotracker red positive vacuoles per plasmatocyte. *n*=21 and 20, respectively. Statistical analysis carried out via Mann-Whitney test, * represents p<0.05. (E) scatterplot showing proportion of vacuoles counted which were lysotracker red positive (i.e., acidified). *n*=21 and 20, respectively. Statistical analyses carried out via unpaired *t*-tests, **** represents p<0.0001.

### Ecdysone signalling contributes to regulation of subpopulation identity

In the context of *Drosophila* development and metamorphosis, the levels of the steroid hormone ecdysone (20-hydroxyecdysone) are absolutely central to a multitude of developmental events (Nicolson et al., 2015) – including the switching of plasmatocytes between different behavioural and transcriptional states (Regan et al., 2013; Sampson et al., 2013) and the ability of embryonic plasmatocytes to mount an effective immune response to bacterial infection (Tan et al., 2014). Given these important roles in programming of *Drosophila* immune cells and the association of ecdysone with developmental transitions during which there are significant changes in plasmatocyte subpopulation numbers, we sought to investigate whether this hormone also plays an instructive role in establishing subpopulation identity. A dominant-negative isoform of the nuclear ecdysone receptor (*UAS-EcR.B1^ΔC655^;* referred to hereafter as *EcR-DN*; Cherbas et al., 2003) was therefore specifically expressed in all plasmatocytes via *srpHemo-GAL4* and *crq-GAL4*, with subpopulations labelled via GAL4*-*independent *VT-RFP* reporters (Coates et al., 2021), with the percentage of plasmatocytes within each subpopulation then determined (Fig. 6A-D’).

**Figure 6.**
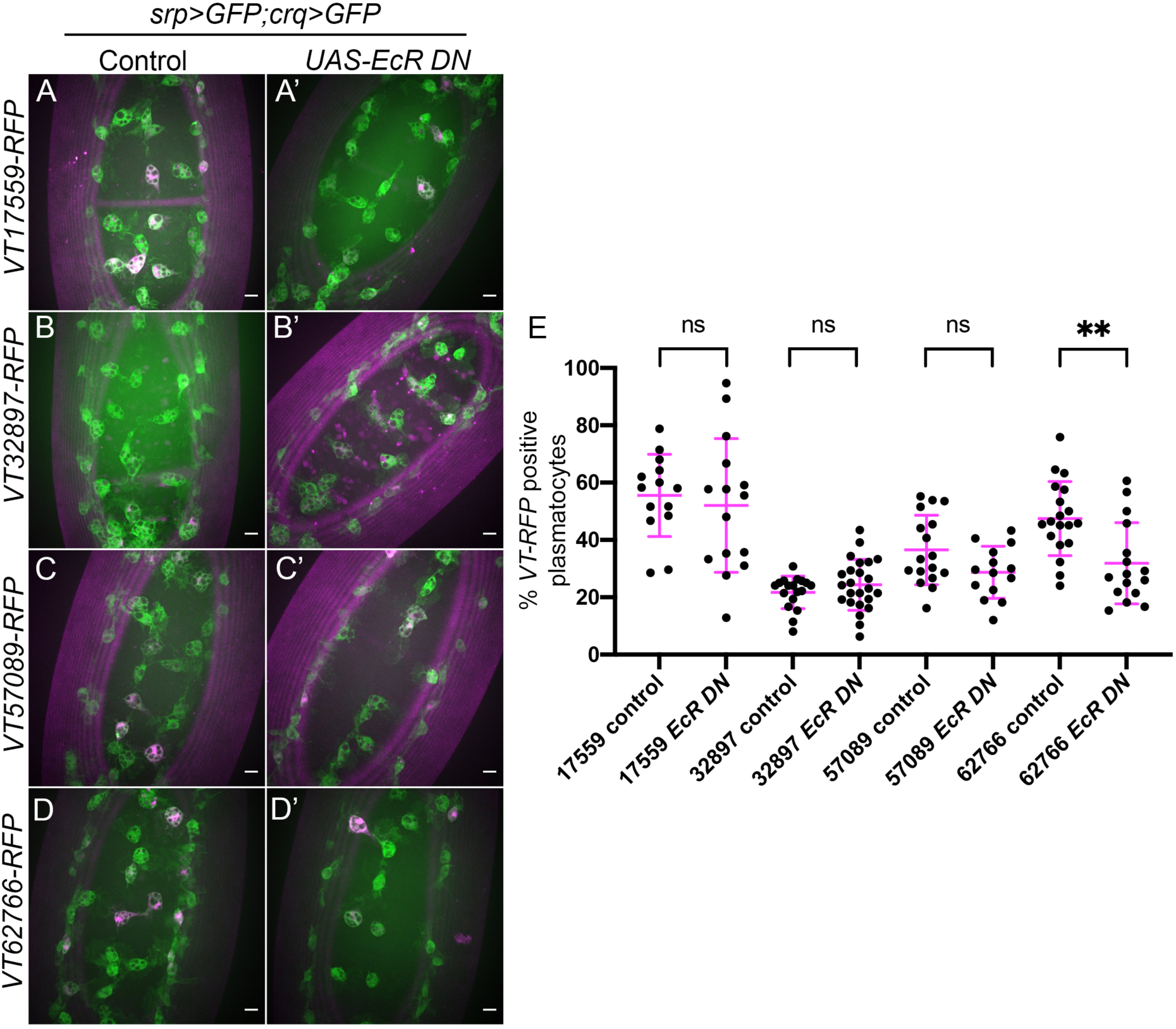
Ecdysone is required for establishing the *VT62766* subpopulation in the embryo. (A-D’) representative maximum projection images of the ventral midline of control embryos (A-D) and embryos expressing *UAS-EcR^ΔC655^* specifically in plasmatocytes (A’-D’) at stage 15. Plasmatocytes labelled via *srpHemo-GAL4,UAS-GFP* and *crq-GAL4,UAS-GFP* (green) while subpopulation plasmatocytes are labelled via *VT-RFP* (magenta). Anterior is up in all images, scale bars denote 10µm. (E) Scatterplot showing proportions of plasmatocytes within subpopulations in the presence and absence of pan- plasmatocyte *UAS-EcR^ΔC655^* expression. *n*=14, 15, 18, 24, 17, 14, 19 and 16, respectively. Statistical analyses carried out via unpaired t- tests, ** represents p<.0.01.

The proportion of plasmatocytes within the *VT17559* and *VT32897* subpopulations was unchanged on expression of EcR-DN, suggesting that ecdysone signalling is not responsible for establishing these subpopulations in the embryo (Fig. 6E). Although the *VT57089* subpopulation exhibited a trend suggesting a modest decrease in numbers in the presence of EcR-DN, this was not statistically significant (Fig. 6E). By contrast, the *VT62766* subpopulation exhibited a 30% decrease of subpopulation plasmatocytes in the presence of EcR-DN, suggesting ecdysone signalling is autonomously required in plasmatocytes for the differentiation and/or maintenance of this subpopulation in the embryo (Fig. 6E).

*VT62766*-labelled cells appear absent during late larval stages but can once more be found in large numbers in pupae (Coates et al., 2021). Hemocytes are exposed to multiple waves of ecdysone during pupal development. Therefore, to test whether ecdysone signalling contributed to re-emergence of this population of cells, we manipulated ecdysone signalling specifically within haemocytes in pupae. As in the embryo, blocking ecdysone signalling within the majority of plasmatocytes (using *hml(Δ)-GAL4* to drive expression from *UAS-GFP* and *UAS-EcR-DN*) decreased the numbers of cells that could be labelled via the *VT62766-RFP* reporter (Fig. 7). Clear morphological differences were also obvious comparing plasmatocytes in the thorax in controls and upon overexpression of EcR-DN (Fig. 7A-B).

**Figure 7.**
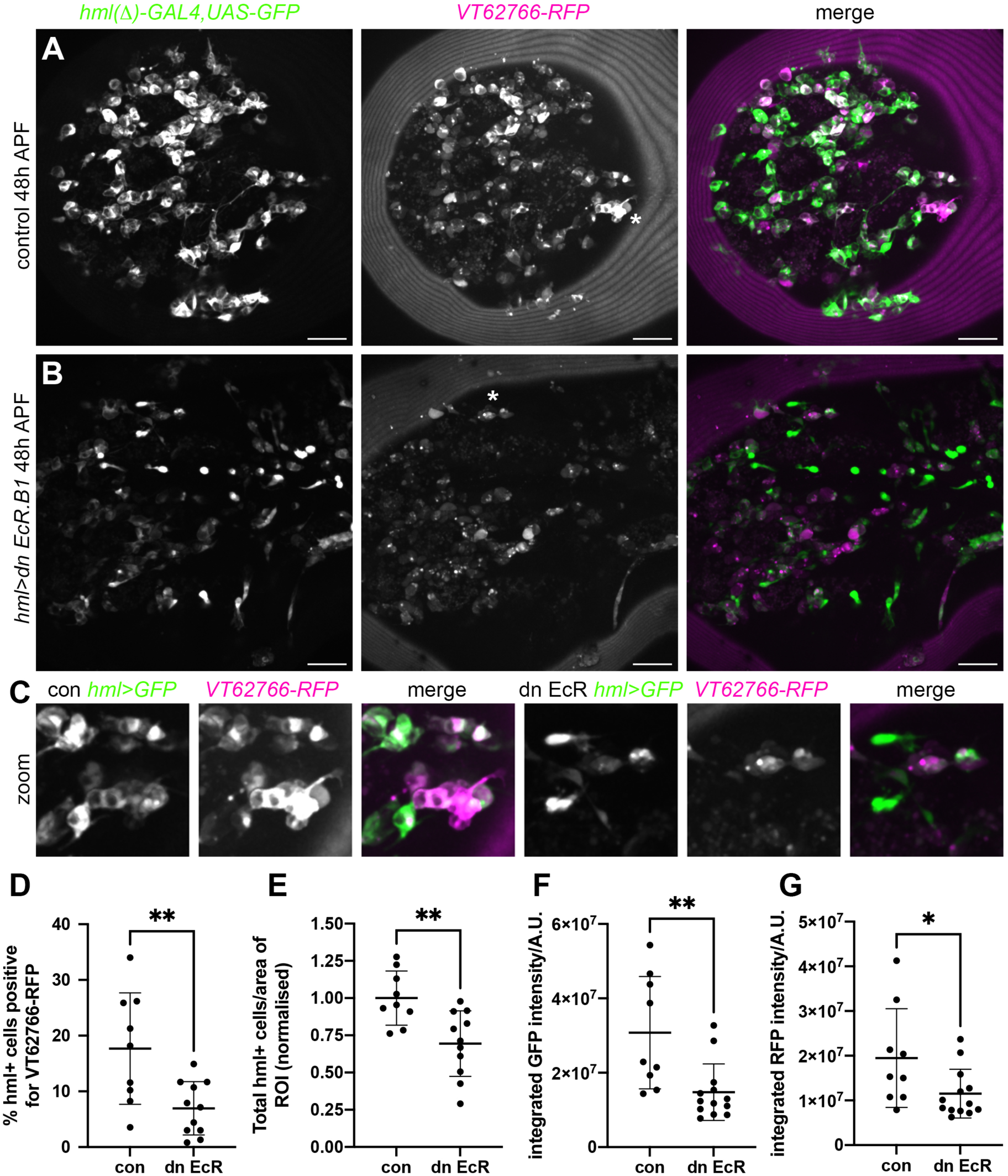
Ecdysone signalling regulates *VT62766* subpopulation cells in pupae. (A-B) Maximum projections of thoracic regions at 48h APF of a control pupa (A) and pupa in which dominant negative EcR.B1 (*hml>dn EcR.B1*, B) was expressed in hemocytes via *hml(D)- GAL4*. Pupae contain *hml(Δ)-GAL4,UAS-GFP* (*hml>GFP*, green in merge) and *VT62766-RFP* (magenta in merge) to label hemocytes and *VT62766*-expressing cells, respectively. (C) zooms of asterisked areas in (A) and (B), which contain examples of hemocytes positive for both GFP and RFP. Scale bars denote 50µm (A-B). (D-F) Scatterplots showing percentage of *hml*-positive cells that also express *VT62766-RFP* (D), total number of *hml*-positive cells per area of the thorax analysed (normalised according to the control mean) and quantification of RFP levels within *hml*-positive cells (integrated RFP fluorescent intensity under a GFP mask; F). Lines and error bars represent mean and standard deviation; * and ** denote p<0.05 and p<0.01, respectively, following Student’s t-tests (D-E) or Mann-Whitney test (F); n=9 and 11 pupae for control and dn EcR pupae.

Overall, our data provide further evidence that the immune system of *Drosophila* comprises of heterogeneous subpopulations of plasmatocytes, akin to vertebrate macrophages. The identities of plasmatocyte subpopulations appear to be modulated by distinct processes, and we have identified signalling pathways (Simu and ecdysone signalling) involved in the establishment of specific subpopulations.

## Discussion

We and others have previously demonstrated macrophage heterogeneity in *Drosophila* (Cattenoz et al., 2020; Cho et al., 2020; Coates et al., 2021; Fu et al., 2020; Tattikota et al., 2020). In this study we investigated mechanisms regulating *Drosophila* macrophage subpopulations *in vivo*, focusing on signalling pathways associated with ingestion of apoptotic cells and developmental transitions and using variations in subpopulation numbers induced by apoptotic cell challenge and developmental stage as experimental tools (Coates et al., 2021). We show that the phosphatidylserine receptor Simu antagonises *VT57089* subpopulation fate and is necessary for reprogramming events that reduce numbers of this subpopulation in the face of excess apoptotic cells. Consistent with a role downstream of Simu, the calcium-permeable cation channel Amo also regulated this subpopulation. Amo was implicated in mediating effective phagosome acidification, which may represent an important process in modulating numbers of cells within the *VT57089* subpopulation. Interestingly, Simu-dependent efferocytosis did not affect other subpopulation identities, while other apoptotic cell receptors (Crq and Drpr) did not seem to integrate apoptotic cell sensing and reprogramming of immune cells. Finally, we show that ecdysone signalling, itself associated with developmental timepoints featuring significant alternations in subpopulation numbers, also impacts identity of specific subpopulations. Taken together, individual plasmatocyte subpopulations are regulated by distinct processes at specific developmental stages. These results further reinforce the model that plasmatocytes exist as a heterogeneous population of cells that are programmed (and/or re-programmed) in response to the precise *in vivo* microenvironment.

*Repo* mutants have previously been used in genetic approaches to challenge plasmatocytes with large amounts of apoptosis (Armitage et al., 2020; Coates et al., 2021; Raymond et al., 2022). Loss of *repo* prevents the specification of glia, another important phagocyte lineage in the embryo (Xiong et al., 1994). This results in increased numbers of uncleared apoptotic cells, as fewer phagocytes remain to clear dying cells (Shklyar et al., 2014). The resulting challenge impairs effective migration and wound responses (Armitage et al., 2020), and alters subpopulation numbers (Coates et al., 2021). Plasmatocytes within *repo* mutant embryos maintain expression of pan-plasmatocyte reporters (e.g., *srpHemo-GAL4*, *crq-GAL4, pxn-GAL4*) and are able to efficiently phagocytose apoptotic cells (Armitage et al., 2020). Nonetheless, we cannot exclude the possibility that glia can also influence plasmatocyte specification independently of apoptosis.

Here we demonstrate a role via which Simu can antagonise acquisition of specific subpopulation identities. In both *repo* and *simu* mutant embryos, plasmatocytes face elevated levels of apoptosis (Armitage et al., 2020; Roddie et al., 2019). However, increasing the levels of apoptotic cells triggered by loss of *repo* has a broad effect on the expression of multiple subpopulation reporters, whereas loss of *simu* is more specific. This suggests that, in some circumstances, contact with apoptotic cells may not be sufficient for changes in subpopulation identity. That loss of *simu* function can block *repo*-induced changes in reporter expression suggests that signalling through this apoptotic cell clearance receptor is mediating these alterations in plasmatocyte identity. Consistent with a role for Simu-dependent signalling, a cation channel known to operate downstream of Simu, Amo (Van Goethem et al., 2012), also impacts subpopulation numbers. However, the precise mechanisms remain to be determined, not least since Simu lacks an intracellular signalling domain (Kurant et al., 2008).

The human homolog of *amo* is *PKD2*, which encodes Polycystin-2 (PC-2). Mutations in *PKD2* and *PKD1* cause autosomal dominant polycystic kidney disease (ADPKD), a relatively common genetic nephropathy, wherein tubular epithelial cells proliferate to form cysts, ultimately resulting in renal failure (Ong & Harris, 2015). Though the exact mechanisms involved in the pathogenesis of ADPKD remain poorly understood, there appears to be a role for immune cells in cyst expansion, with macrophage polarisation also implicated. Monocyte chemoattractant protein (MCP-1) and macrophage migration inhibitory factor (MIF) lead to an initial influx of pro-inflammatory macrophages, which then polarise towards a more anti-inflammatory, pro-proliferative activation state, driving cyst expansion and disease progression (Cassini et al., 2018; Chen et al., 2015; Karihaloo et al., 2011). Our data revealed decreased phagosome acidification in *amo;repo* mutant plasmatocytes compared to *repo* only controls. The changes in the type of innate immune cells present in ADPKD and in the fly embryo on loss of *PKD2*/*amo* suggests manipulation of phagosome maturation may represent a novel pathway to target with respect to further understanding pathogenesis of the disease.

Amo is a cation channel and calcium signals have long been linked to phagocytic events, albeit the data has not always proven consistent (Westlake et al., 2019). A recent paper beautifully delineates a requirement of calcium nanodomains for activation of dynamin during phagocytosis (Davis et al., 2020), though the key channels here were NAADP-regulated two-pore channels. Phagocytosis is also associated with more global elevations in cytoplasmic calcium and these authors speculate that these may regulate changes in mitochondrial energetics and gene expression. It is plausible that Amo might contribute to global Ca^2+^ signals under conditions of phagocytic stress. In turn, this may facilitate a contribution to changes in gene expression necessary to reprogramme macrophage subpopulations. Notably, calcium signalling is already known to be associated with changes in gene expression following apoptotic cell clearance in *Drosophila* (Weavers et al., 2016).

The steroid hormone ecdysone is known to be involved in mediating large-scale phenotypic changes in the overall plasmatocyte population associated with different developmental stages (Jiang et al., 1997; Regan et al., 2013; Sampson et al., 2013; Tan et al., 2014). For example, larval plasmatocytes, which are typically sessile and proliferative, become highly migratory and phagocytic at the onset of metamorphosis in response to ecdysone (Regan et al., 2013). Similarly, embryonic plasmatocytes are unable to mount an effective immune response until being exposed to ecdysone at stage 12 (Tan et al., 2014). Expression of a dominant-negative ecdysone receptor isoform revealed a decreased proportion of plasmatocytes within the *VT62766* subpopulation in both embryos and pupae. It is therefore possible that ecdysone is required to help establish this plasmatocyte subpopulation. Other subpopulations were not affected by this approach, highlighting specificity in how subpopulation identity is established and controlled.

Ecdysone signalling has been shown to drive expression of phagocytic genes in pupae (Regan et al., 2013), stimulate motility and cytoplasmic rearrangements (Edwards et al., 2018; Sampson et al., 2013), and help establish immune responses (Tan et al., 2014). The *VT62766* subpopulation exhibits enhanced wound responses, but reduced rates of phagocytosis (Coates et al., 2021), so while ecdysone could be argued to drive cells towards a more activated state, associated changes in behaviour do not completely align. However, it is important to note that earlier papers quantify behaviour across the total population of plasmatocytes so subpopulation-specific effects could be obscured.

Like steroid hormone signalling, apoptotic cell death is also associated with reprogramming of vertebrate macrophages (Serhan & Savill, 2005). Pro-inflammatory cytokines are inhibited in macrophages following phagocytosis of apoptotic cells (Fadok et al., 1998). Contact with large numbers of apoptotic cells reprograms plasmatocytes away from identities that exhibit less efficient apoptotic cell clearance (Coates et al., 2021), which potentially might signify a less pro-inflammatory state. Loss of an apoptotic cell clearance receptor blocks that effect for at least one discrete population of cells, reinforcing differences between the cells marked using our transgenic reporters.

In summary, we have identified new molecular players involved in determining the acquisition of specific plasmatocyte subpopulation identities in *Drosophila*: Simu and the downstream effector Amo regulate *VT57089* identity, with phagosome acidification a potential point of integration. Meanwhile, ecdysone appears important in establishing identity of *VT62766*- labelled cells in both the embryo and pupa. The role of steroid hormone signalling and apoptosis suggest that mechanisms controlling innate immune cell behaviour in *Drosophila* and vertebrates may be more similar than previously thought. Finally, this further supports the existence of macrophage heterogeneity within this important immune model and will enable use of the fly to further dissect regulation of this important facet of biology *in vivo*.

## Author contributions

EB and IE performed all experimental work and produced the first draft of the paper. EB, IE and MZ analysed the data. EB, AO, IE acquired funding to support the project (DiMeN PhD studentship awarded to EB, IE, and AO; Wellcome/Royal Society Sir Henry Dale Fellowship awarded to IE). All authors contributed to experimental design and edited the manuscript. IE and AO supervised EB as part of his PhD studies with additional training and feedback from MZ.

## Acknowledgements

This work was funded by an MRC Discovery Medicine North (DiMeN DTP) PhD studentship awarded to EB, AO and IE (MR/N013840/1) and a Wellcome/Royal Society Sir Henry Dale Fellowship awarded to IE (102503/Z /13/Z). We thank Emma Bristow for her assistance in the preparation of *Drosophila* pupae for live imaging (Fig. 7) and Juliette Howarth for help with the ecdysone signalling experiments in the embryo. We acknowledge Karen Plant and the University of Sheffield Fly Facility for support with fly husbandry. We are grateful for access to the microscopes of the University of Sheffield Wolfson Light Microscopy Facility and to Darren Robinson and Nicholas Van Hateren for assistance with imaging. This work used a Perkin Elmer spinning disk system (MRC grant G0700091 and Wellcome grant 077544/Z/05/Z) and a Nikon W1 spinning disk system (BBSRC ALERT2021 award BB/V019368/1) housed in the Wolfson LMF. This work would not be possible without reagents and resources obtained from or maintained by the Bloomington *Drosophila* Stock Centre (NIH P40OD018537) and Flybase (MRC grant MR/N030117/1). We thank Brian Stramer, Estee Kurant and Daria Siekhaus for providing additional fly lines.

## References

Armitage, E. L., Roddie, H. G., & Evans, I. R. (2020). Overexposure to apoptosis via disrupted glial specification perturbs Drosophila macrophage function and reveals roles of the CNS during injury. Cell Death and Disease. 10.1038/s41419-020-02875-2

Barolo, S., Carver, L. A., & Posakony, J. W. (2000). GFP and β-galactosidase transformation vectors for promoter/enhancer analysis in Drosophila. In BioTechniques (Vol. 29, Issue 4, pp. 726–732). Eaton Publishing Company. 10.2144/00294bm10

Brückner, K., Kockel, L., Duchek, P., Luque, C. M., Rørth, P., & Perrimon, N. (2004). The PDGF/VEGF receptor controls blood cell survival in Drosophila. Developmental Cell, 7(1), 73–84. 10.1016/j.devcel.2004.06.007

Cassini, M. F., Kakade, V. R., Kurtz, E., Sulkowski, P., Glazer, P., Torres, R., Somlo, S., & Cantley, L. G. (2018). Mcp1 Promotes Macrophage-Dependent Cyst Expansion in Autosomal Dominant Polycystic Kidney Disease. Journal of the American Society of Nephrology, 29(10), 2471–2481. 10.1681/ASN.2018050518

Cattenoz, P. B., Sakr, R., Pavlidaki, A., Delaporte, C., Riba, A., Molina, N., Hariharan, N., Mukherjee, T., & Giangrande, A. (2020). Temporal specificity and heterogeneity of Drosophila immune cells . The EMBO Journal, 39(12), 1–25. 10.15252/embj.2020104486

Chen, L., Zhou, X., Fan, L. X., Yao, Y., Swenson-Fields, K. I., Gadjeva, M., Wallace, D. P., Peters, D. J. M., Yu, A., Grantham, J. J., & Li, X. (2015). Macrophage migration inhibitory factor promotes cyst growth in polycystic kidney disease. Journal of Clinical Investigation, 125(6), 2399–2412. 10.1172/JCI80467

Cherbas, L., Hu, X., Zhimulev, I., Belyaeva, E., & Cherbas, P. (2003). EcR isoforms in Drosophila: testing tissue-specific requirements by targeted blockade and rescue. Development, 130(2), 271–284. 10.1242/DEV.00205

Cho, B., Yoon, S. H., Lee, D., Koranteng, F., Tattikota, S. G., Cha, N., Shin, M., Do, H., Hu, Y., Oh, S. Y., Lee, D., Vipin Menon, A., Moon, S. J., Perrimon, N., Nam, J. W., & Shim, J. (2020). Single-cell transcriptome maps of myeloid blood cell lineages in Drosophila. Nature Communications, 11(1). 10.1038/s41467-020-18135-y

Coates, J. A., Brooks, E., Brittle, A. L., Armitage, E. L., Zeidler, M. P., & Evans, I. R. (2021). Identification of functionally distinct macrophage subpopulations in drosophila. ELife, 10. 10.7554/ELIFE.58686

Cornwell, W. D., Kim, V., Fan, X., Vega, M. E., Ramsey, F. V., Criner, G. J., & Rogers, T. J. (2018). Activation and polarization of circulating monocytes in severe chronic obstructive pulmonary disease. BMC Pulmonary Medicine, 18(1). 10.1186/S12890-018-0664-Y

Davis, L. C., Morgan, A. J., & Galione, A. (2020). NAADP -regulated two-pore channels drive phagocytosis through endo-lysosomal Ca 2+ nanodomains, calcineurin and dynamin . The EMBO Journal, 39(14). 10.15252/embj.2019104058

de Gaetano, M., Crean, D., Barry, M., & Belton, O. (2016). M1- and M2-type macrophage responses are predictive of adverse outcomes in human atherosclerosis. Frontiers in Immunology, 7(JUL), 275. 10.3389/FIMMU.2016.00275/BIBTEX

Edwards, S. S., Delgado, M. G., Nader, G. P. de F., Piel, M., Bellaïche, Y., Lennon-Duménil, A. M., & Glavic, Á. (2018). An in vitro method for studying subcellular rearrangements during cell polarization in Drosophila melanogaster hemocytes. Mechanisms of Development, 154, 277–286. 10.1016/j.mod.2018.08.003

Evans, Iwan R., Hu, N., Skaer, H., & Wood, W. (2010). Interdependence of macrophage migration and ventral nerve cord development in Drosophila embryos. *Development (Cambridge*, England), 137(10), 1625–1633. 10.1242/dev.046797

Evans, Iwan Robert, Rodrigues, F. S. L. M., Armitage, E. L., & Wood, W. (2015). Draper/CED-1 Mediates an Ancient Damage Response to Control Inflammatory Blood Cell Migration In Vivo. Current Biology : CB, 25(12), 1606–1612. 10.1016/j.cub.2015.04.037

Fadok, V. A., Bratton, D. L., Konowal, A., Freed, P. W., Westcott, J. Y., & Henson, P. M. (1998). Macrophages That Have Ingested Apoptotic Cells In Vitro Inhibit Proinflammatory Cytokine Production Through Autocrine/Paracrine Mechanisms Involving TGF-, PGE2, and PAF. J. Clin. Invest, 101(4), 890–898.

Franc, N. C., Dimarcq, J.-L., Lagueux, M., Hoffmann, J., & Ezekowitz, R. A. B. (1996). Croquemort, A Novel Drosophila Hemocyte/Macrophage Receptor that Recognizes Apoptotic Cells. Immunity, 4(5), 431–443. 10.1016/S1074-7613(00)80410-0

Freeman, M. R., Delrow, J., Kim, J., Johnson, E., & Doe, C. Q. (2003). Unwrapping Glial Biology: Gcm Target Genes Regulating Glial Development, Diversification, and Function brain barrier that isolates and protects neural tissue. Glia mediate many brain responses to injury and neuro-degenerative diseases (Wyss-Coray and Muck. In Neuron (Vol. 38).

Fu, Y., Huang, X., Zhang, P., van de Leemput, J., & Han, Z. (2020). Single-cell RNA sequencing identifies novel cell types in Drosophila blood. Journal of Genetics and Genomics, 47(4), 175–186. 10.1016/j.jgg.2020.02.004

Gordon, S., & Plüddemann, A. (2017). Tissue macrophages: heterogeneity and functions. BMC Biology, 15(1), 53. 10.1186/s12915-017-0392-4

Guillou, A., Troha, K., Wang, H., Franc, N. C., & Buchon, N. (2016). The Drosophila CD36 Homologue croquemort Is Required to Maintain Immune and Gut Homeostasis during Development and Aging. PLoS Pathogens, 12(10). 10.1371/journal.ppat.1005961

Gyoergy, A., Roblek, M., Ratheesh, A., Valoskova, K., Belyaeva, V., Wachner, S., Matsubayashi, Y., Sánchez-Sánchez, B. J., Stramer, B., & Siekhaus, D. E. (2018). Tools Allowing Independent Visualization and Genetic Manipulation of *Drosophila melanogaster* Macrophages and Surrounding Tissues. G3 Genes |Genomes|Genetics, 8(3), 845–857. 10.1534/g3.117.300452

Halter, D. a, Urban, J., Rickert, C., Ner, S. S., Ito, K., Travers, a a, & Technau, G. M. (1995). The homeobox gene repo is required for the differentiation and maintenance of glia function in the embryonic nervous system of Drosophila melanogaster. *Development (Cambridge*, England), 121(2), 317–332.

Hume, D. A. (2015). The many alternative faces of macrophage activation. Frontiers in Immunology, 6(JUL), 370. 10.3389/FIMMU.2015.00370/BIBTEX

Jiang, C., Baehrecke, E. H., & Thummel, C. S. (1997). Steroid regulated programmed cell death during Drosophila metamorphosis. Development, 124(22), 4673–4683. 10.1242/DEV.124.22.4673

Karihaloo, A., Koraishy, F., Huen, S. C., Lee, Y., Merrick, D., Caplan, M. J., Somlo, S., & Cantley, L. G. (2011). Macrophages promote cyst growth in polycystic kidney disease. Journal of the American Society of Nephrology : JASN, 22(10), 1809–1814. 10.1681/ASN.2011010084

Kinchen, J. M., & Ravichandran, K. S. (2008). Phagosome maturation: going through the acid test. Nature Reviews. Molecular Cell Biology, 9(10), 781–795. 10.1038/nrm2515

Kurant, E., Axelrod, S., Leaman, D., & Gaul, U. (2008). Six-Microns-Under Acts Upstream of Draper in the Glial Phagocytosis of Apoptotic Neurons. Cell, 133(3), 498–509. 10.1016/j.cell.2008.02.052

Kvon, E. Z., Kazmar, T., Stampfel, G., Yáñez-Cuna, J. O., Pagani, M., Schernhuber, K., Dickson, B. J., & Stark, A. (2014). Genome-scale functional characterization of Drosophila developmental enhancers in vivo. Nature. 10.1038/nature13395

Lebestky, T., Chang, T., Hartenstein, V., & Banerjee, U. (2000). Specification of Drosophila Hematopoietic Lineage by Conserved Transcription Factors. Science, 288(5463), 146–149. 10.1126/science.288.5463.146

Leitão, A. B., & Sucena, É. (2015). Drosophila sessile hemocyte clusters are true hematopoietic tissues that regulate larval blood cell differentiation. ELife, 2015(4), 1–38. 10.7554/eLife.06166

Logan, M. A., Hackett, R., Doherty, J., Sheehan, A., Speese, S. D., & Freeman, M. R. (2012). Negative regulation of glial engulfment activity by Draper terminates glial responses to axon injury. Nature Neuroscience, 15(5), 722–730. 10.1038/nn.3066

Makhijani, K., Alexander, B., Rao, D., Petraki, S., Herboso, L., Kukar, K., Batool, I., Wachner, S., Gold, K. S., Wong, C., O’Connor, M. B., & Brückner, K. (2017). Regulation of Drosophila hematopoietic sites by Activin-β from active sensory neurons. Nature Communications 2017 8:1, 8(1), 1–12. 10.1038/ncomms15990

Manaka, J., Kuraishi, T., Shiratsuchi, A., Nakai, Y., Higashida, H., Henson, P., & Nakanishi, Y. (2004). Draper-mediated and Phosphatidylserine-independent Phagocytosis of Apoptotic Cells by Drosophila Hemocytes/Macrophages. Journal of Biological Chemistry, 279(46), 48466–48476. 10.1074/jbc.M408597200

Matsubayashi, Y., Louani, A., Dragu, A., Sánchez-Sánchez, B. J., Serna-Morales, E., Yolland, L., Gyoergy, A., Vizcay, G., Fleck, R. A., Heddleston, J. M., Chew, T.-L., Siekhaus, D. E., & Stramer, B. M. (2017). A Moving Source of Matrix Components Is Essential for De Novo Basement Membrane Formation. Current Biology, 27(22), 3526–3534.e4. 10.1016/j.cub.2017.10.001

Murray, P. J. (2017). Macrophage Polarization. Annual Review of Physiology, 79(1), 541–566. 10.1146/annurev-physiol-022516-034339

Nicolson, S., Denton, D., & Kumar, S. (2015). Ecdysone-mediated programmed cell death in Drosophila. The International Journal of Developmental Biology, 59(1-2–3), 23–32. 10.1387/ijdb.150055sk

Ong, A. C. M., & Harris, P. C. (2015). A polycystin-centric view of cyst formation and disease: the polycystins revisited. Kidney International, 88(4), 699–710. 10.1038/ki.2015.207

Orecchioni, M., Ghosheh, Y., Pramod, A. B., & Ley, K. (2019). Macrophage Polarization: Different Gene Signatures in M1(LPS+) vs. Classically and M2(LPS–) vs. Alternatively Activated Macrophages. Frontiers in Immunology, 0(MAY), 1084. 10.3389/FIMMU.2019.01084

Page, D. T., & Olofsson, B. (2008). Multiple roles for apoptosis facilitating condensation of the Drosophila ventral nerve cord. Genesis, 46(2), 61–68. 10.1002/dvg.20365

Pfeiffer, B. D., Ngo, T. T. B., Hibbard, K. L., Murphy, C., Jenett, A., Truman, J. W., & Rubin, G. M. (2010). Refinement of tools for targeted gene expression in Drosophila. Genetics. 10.1534/genetics.110.119917

Raymond, M. H., Davidson, A. J., Shen, Y., Tudor, D. R., Lucas, C. D., Morioka, S., Perry, J. S. A., Krapivkina, J., Perrais, D., Schumacher, L. J., Campbell, R. E., Wood, W., & Ravichandran, K. S. (2022). Live cell tracking of macrophage efferocytosis during *Drosophila* embryo development in vivo. Science, 375(6585), 1182–1187. 10.1126/SCIENCE.ABL4430

Regan, J. C., Brandão, A. S., Leitão, A. B., Mantas Dias, Â. R., Sucena, É., Jacinto, A., & Zaidman-Rémy, A. (2013). Steroid Hormone Signaling Is Essential to Regulate Innate Immune Cells and Fight Bacterial Infection in Drosophila. PLoS Pathogens, 9(10), e1003720. 10.1371/journal.ppat.1003720

Roddie, H. G., Armitage, E. L., Coates, J. A., Johnston, S. A., & Evans, I. R. (2019). Simu-dependent clearance of dying cells regulates macrophage function and inflammation resolution. PLOS Biology, 17(5), e2006741. 10.1371/journal.pbio.2006741

Sampson, C. J., Amin, U., & Couso, J. P. (2013). Activation of Drosophila hemocyte motility by the ecdysone hormone. Biology Open, 2(12), 1412–1420. 10.1242/bio.20136619

Schindelin, J., Arganda-Carreras, I., Frise, E., Kaynig, V., Longair, M., Pietzsch, T., Preibisch, S., Rueden, C., Saalfeld, S., Schmid, B., Tinevez, J. Y., White, D. J., Hartenstein, V., Eliceiri, K., Tomancak, P., & Cardona, A. (2012). Fiji: An open-source platform for biological-image analysis. Nature Methods, 9(7), 676–682. 10.1038/nmeth.2019

Serhan, C. N., & Savill, J. (2005). Resolution of inflammation: the beginning programs the end. Nature Immunology, 6(12), 1191–1197. 10.1038/ni1276

Shaykhiev, R., Krause, A., Salit, J., Strulovici-Barel, Y., Harvey, B.-G., O’Connor, T. P., & Crystal, R. G. (2009). Smoking-dependent reprogramming of alveolar macrophage polarization: implication for pathogenesis of chronic obstructive pulmonary disease. Journal of Immunology (Baltimore, Md. : 1950), 183(4), 2867–2883. 10.4049/JIMMUNOL.0900473

Shklyar, B., Sellman, Y., Shklover, J., Mishnaevski, K., Levy-Adam, F., & Kurant, E. (2014). Developmental regulation of glial cell phagocytic function during Drosophila embryogenesis. Developmental Biology, 393(2), 255–269. 10.1016/j.ydbio.2014.07.005

Sinenko, S. A., & Mathey-Prevot, B. (2004). Increased expression of Drosophila tetraspanin, Tsp68C, suppresses the abnormal proliferation of ytr-deficient and Ras/Raf-activated hemocytes. Oncogene, 23(56), 9120–9128. 10.1038/sj.onc.1208156

Sonnenfeld, M. J., & Jacobs, J. R. (1995). Macrophages and glia participate in the removal of apoptotic neurons from theDrosophila embryonic nervous system. The Journal of Comparative Neurology, 359(4), 644–652. 10.1002/cne.903590410

Stramer, B., Wood, W., Galko, M. J., Redd, M. J., Jacinto, A., Parkhurst, S. M., & Martin, P. (2005). Live imaging of wound inflammation in Drosophila embryos reveals key roles for small GTPases during in vivo cell migration. Journal of Cell Biology, 168(4), 567–573. 10.1083/jcb.200405120

Tan, K. L., Vlisidou, I., & Wood, W. (2014). Ecdysone mediates the development of immunity in the drosophila embryo. Current Biology, 24(10), 1145–1152. 10.1016/j.cub.2014.03.062

Tattikota, S. G., Cho, B., Liu, Y., Hu, Y., Barrera, V., Steinbaugh, M. J., Yoon, S. H., Comjean, A., Li, F., Dervis, F., Hung, R. J., Nam, J. W., Sui, S. H., Shim, J., & Perrimon, N. (2020). A single-cell survey of drosophila blood. ELife, 9, 1–35. 10.7554/eLife.54818

Van Goethem, E., Silva, E. A., Xiao, H., & Franc, N. C. (2012). The Drosophila TRPP cation channel, PKD2 and Dmel/Ced-12 act in genetically distinct pathways during apoptotic cell clearance. PloS One, 7(2), e31488. 10.1371/journal.pone.0031488

Watnick, T. J., Jin, Y., Matunis, E., Kernan, M. J., & Montell, C. (2003). A Flagellar Polycystin-2 Homolog Required for Male Fertility in Drosophila. Current Biology, 13(24), 2179–2184. 10.1016/J.CUB.2003.12.002

Weavers, H., Evans, I. R., Martin, P., & Wood, W. (2016). Corpse Engulfment Generates a Molecular Memory that Primes the Macrophage Inflammatory Response. Cell, 165(7), 1658–1671. 10.1016/j.cell.2016.04.049

Westman, J., Grinstein, S., & Maxson, M. E. (2019). Revisiting the role of calcium in phagosome formation and maturation. Journal of Leukocyte Biology, 106(4), 837–851. 10.1002/JLB.MR1118-444R

Wood, W., & Martin, P. (2017). Macrophage Functions in Tissue Patterning and Disease: New Insights from the Fly. Developmental Cell, 40(3), 221–233. 10.1016/j.devcel.2017.01.001

Xiong, W. C., Okano, H., Patel, N. H., Blendy, J. A., & Montell, C. (1994). repo encodes a glial-specific homeo domain protein required in the Drosophila nervous system. Genes and Development. 10.1101/gad.8.8.981

Zanet, J., Stramer, B., Millard, T., Martin, P., Payre, F., & Plaza, S. (2009). Fascin is required for blood cell migration during Drosophila embryogenesis. Development, 136(15), 2557–2565. 10.1242/dev.036517

Zirin, J., Cheng, D., Dhanyasi, N., Cho, J., Dura, J. M., VijayRaghavan, K., & Perrimon, N. (2013). Ecdysone signaling at metamorphosis triggers apoptosis of Drosophila abdominal muscles. Developmental Biology, 383(2), 275–284. 10.1016/J.YDBIO.2013.08.029

